# Genetic modification of primary airway epithelial cell cultures defines MX1 as a barrier for zoonotic influenza A viruses in the human respiratory epithelium

**DOI:** 10.64898/2026.10.19.759965

**Authors:** Karen Stevers, Blandina I. Oliveira Esteves, Étori Aguiar Moreira, Sarah Jordi, Laura Laloli, Cinzia Moscardelli, Manon F. Licheri, Dan Wettstein, Ada Mescoli, Mike Mwanga, Alina T. Baltensperger, Larise Oberholster, Philip V’kovski, Gert Zimmer, Ronald Dijkman

**Affiliations:** Institute for Infectious Diseases, University of Bern, Bern, Switzerland; Graduate School for Cellular and Biomedical Sciences, University of Bern, Bern, Switzerland; Institute of Virology and Immunology IVI, Mittelhäusern, Switzerland; Department of Infectious Diseases and Pathobiology, Vetsuisse Faculty, University of Bern, Bern, Switzerland; Multidisciplinary Center for Infectious Diseases, University of Bern, Bern, Switzerland

## Abstract

The antiviral innate immune response is a major species barrier that emerging viruses must overcome to infect and transmit efficiently in new hosts. In humans, the interferon-stimulated *MX1* gene encodes the antiviral MX1 protein, which restricts zoonotic influenza A viruses (IAVs). Resistance to MX1 is considered a key feature of human-adapted IAVs and a potential determinant of zoonotic and pandemic risk. However, MX1-mediated restriction and viral escape have mostly been investigated in cell lines and animal models, which incompletely recapitulate virus–host interactions in the human airway epithelium, representing the primary site of IAV replication. Although primary human airway epithelial cell (hAEC) cultures, such as airway organoids and air-liquid interface (ALI) cultures, provide a physiologically relevant model, their use in studying MX1 restriction has been limited by challenges in achieving efficient and reliable genetic modification.

Here, we used engineered virus-like particles (eVLPs) to deliver Cas9-sgRNA ribonucleoprotein complexes into primary hAECs and disrupt MX1 expression. This approach generated MX1-knockout airway organoids and organoid-derived hAEC ALI cultures with >90% genomic-level editing efficiency and near-complete loss of MX1 protein, while preserving epithelial differentiation at the air-liquid interface. MX1 knockout in hAEC ALI cultures had only a minor effect on the replication kinetics of the human-adapted H1N1pdm09 strain. By contrast, MX1 knockout markedly impaired interferon lambda-mediated restriction of the zoonotic bovine H5N1/Texas and avian H7N9/Anhui strains, resulting in viral titers up to 3-4 logs TCID_50_ higher than in MX1-competent cultures. These findings provide direct evidence that MX1 restricts zoonotic IAVs in primary human airway epithelium. Our study establishes a biologically relevant platform for assessing the sensitivity of emerging IAVs to a key antiviral host factor and thereby informs evaluations into their zoonotic and pandemic risk.

## Introduction

Since the 1918 influenza pandemic, Influenza A viruses (IAVs) have remained a major global health threat, causing annual seasonal epidemics and sporadic pandemics with substantial morbidity and mortality worldwide (1). Zoonotic influenza strains, such as those from avian, bovine or swine origin sporadically can cause severe diseases in humans but are usually not transmitted from human to human, limiting their pandemic risk (2,3). However, continued evolution of IAVs through genetic drift or reassortment may lead to viruses that can efficiently spread within the human population with devastating consequences in immune-naïve communities (4). For example, the reassortment of avian, swine, and human viruses led to the 2009 pandemic H1N1 strain, which resulted in a significant burden of disease in the human population (5–7).The continued animal-to-animal transmission of highly pathogenic H5N1 and H7N9 strains further emphasizes the need to evaluate which emerging influenza A viruses pose a significant risk to the human population (8).

To productively infect humans, zoonotic IAVs must overcome several barriers that limit infection and replication in the human respiratory tract. These include receptor availability, the ability of the virus to exploit host factors for replication, and the capacity to evade restriction factors induced by the host’s antiviral innate immune response (9–11). A central component of the antiviral innate immune response is the interferon (IFN) response, which protects cells against invading pathogens by inducing the expression of interferon-stimulated genes (ISGs) (12). One of those ISGs, *MX1,* encodes the myxovirus resistance protein 1 (MX1), and is considered a key determinant of the human species barrier against zoonotic IAVs. This protein inhibits viral replication by sequestering incoming and newly synthesized viral ribonucleoprotein complexes in the cytoplasm (13,14). Experimental studies in cell culture and mouse models have shown that human MX1 strongly restricts zoonotic IAVs, while human-adapted IAVs are largely resistant to its antiviral activity (15–18). This difference in sensitivity between viruses from animal and human origin is thought to depend largely on differences in the viral nucleoprotein and was later attributed to specific amino acids in the surface-facing body domain of the protein (17,19,20). The amino acids associated with MX1-resistance are widespread in human IAV strains but less common in strains circulating in animals, suggesting strong selective pressure to evade MX1 in the human host through adaptive mutations in the viral nucleoprotein (17,21,22). Although common animal reservoirs for IAVs, such as swine and birds, also express Mx proteins, these proteins display reduced antiviral activity compared to their human counterpart (17,23–25). Thus, resistance to MX1 is considered a hallmark of adaptation to the human host and consequently, a determinant of zoonotic potential and pandemic risk. The mechanisms underlying MX1 restriction and viral escape have largely been investigated in immortalized cell lines and transgenic mouse models, which only partially recapitulate virus–host interactions at the primary site of infection, the human airway epithelium. Moreover, these studies relied mainly on MX1 overexpression, which may overestimate the effect of MX1 on IAV replication under physiological conditions (13). Consequently, the exact contribution of MX1 to the inhibition of zoonotic IAVs in a physiologically relevant system remains elusive.

Primary human airway epithelial cell (hAEC) cultures, such as airway organoids (AO) or air-liquid interface (ALI) cultures, are *in vitro* models that closely mimic the architecture and function of the human airway epithelium. They are created by culturing primary airway epithelial cells derived from the nasal, tracheal, or bronchial regions of the respiratory tract in an extracellular matrix (organoids) or on cell culture inserts with the apical side exposed to air (air-liquid interface), which drives differentiation from basal progenitor cells to a pseudostratified epithelium composed of ciliated, secretory, and basal cells (26–28). These models recapitulate cellular diversity, native expression of influenza virus (co-)receptors, and a functional innate immune response (29–32), making them physiologically relevant systems for studying IAV infection. However, generating knockout (KO) hAEC cultures to specifically study the function of species-specific IAV restriction factors, such as MX1, is technically challenging due to the cells’ finite lifespan, rapid senescence, and sensitive differentiation at the air-liquid interface (33). In the past, hAECs have been genetically modified in our laboratory and by others using lentiviral transduction, electroporation of plasmids encoding Cas9 and gRNAs or direct electroporation of the Cas9-sgRNA ribonucleoprotein (RNP) complex (34–38). Drawbacks of these established techniques include limited efficiency, high cytotoxicity, or raised concerns about genomic instability and off-target effects (39–41).

Combining viral delivery with transient RNP expression using engineered virus-like particles (eVLPs) has recently emerged as a new strategy for genome engineering. eVLPs are based on retroviruses such as Murine Moloney Leukemia virus (MMLV) or Human Immunodeficiency Virus (HIV) and package the RNP complex into virions by fusing the Cas9 protein to the retroviral gag-protein (42,43). During virion maturation, the Cas9-Gag fusion protein is cleaved, allowing Cas9 to localize to the nucleus and edit the genome (26,28). eVLPs demonstrated highly efficient editing without associated cytotoxicity or off-target effects in human and mouse intestinal and prostate organoids, primary hematopoietic stem cells and *in vivo* (42,44–47).

To evaluate how MX1 expression in the human airway epithelium affects the replication of IAVs of zoonotic or human origin, we established an eVLP-based genetic-modification pipeline for airway epithelial progenitor cells to generate well-differentiated MX1 knockout (KO) hAEC ALI cultures. eVLP-mediated genome editing overcomes key limitations of existing genome-editing methods, including low efficiency and cytotoxicity, while preserving mucociliary differentiation and epithelial integrity at the air-liquid interface. The resulting well-differentiated MX1 KO hAEC ALI cultures were subsequently infected with avian H7N9, bovine H5N1, and human-adapted H1N1pdm09 influenza A viruses. We found that MX1 knockout significantly reduced the sensitivity of zoonotic bovine H5N1/Texas and avian H7N9/Anhui strains to interferon lambda stimulation, whereas it had only a minor effect on the replication of the human-adapted H1N1pdm09 strain. Altogether, our findings indicate that MX1 expression in the human airway epithelium is a key determinant in the interferon-mediated restriction of zoonotic influenza A viruses.

## Results

### Efficient knock out of MX1 in primary hAECs using eVLPs

We hypothesized that eVLPs could be an attractive tool for genetic modification of primary airway epithelial cells due to the transient nature of the system, minimizing undesired side effects, while simultaneously being highly efficient in a variety of primary cell types. Such high efficiency could reduce the number of passages or subcloning required to create a complete knockout of a gene of interest and preserve the limited lifespan of primary hAECs (33). First, we produced eVLPs by transfecting 293-LTV cells with plasmids encoding the glycoprotein of vesicular stomatitis virus (VSV-G), a MMLV gag-Cas9 fusion protein, a MMLV gag-pro-pol polyprotein making up the retroviral scaffold, and a gRNA targeting *MX1*. We then determined the optimal dose for eVLP transduction by transducing undifferentiated primary bronchial epithelial cells with increasing volumes of supernatant from eVLP producer cells for 24h (Figure 1A). We chose to target undifferentiated hAECs, comprising primary airway epithelial basal stem cells, because prior work indicated that well-differentiated hAEC cultures are not efficiently transduced and have limited self-renewal capacity (37,48). Sanger sequencing, followed by indel quantification at the gRNA target site 72h post-transduction, demonstrated dose-dependent genome editing (Figure 1B). Importantly, when transducing hAECs with the highest doses (eVLPs comprising 50-100% of the total volume), editing efficiencies exceeded 90% at the genomic level (Figure 1B). Furthermore, supplementation with 5 μg/mL of the cationic polymer polybrene during transduction significantly increased editing efficiency (Figure 1B), likely due to enhanced adsorption of viral particles to the cell surface (49).

**Figure 1:**
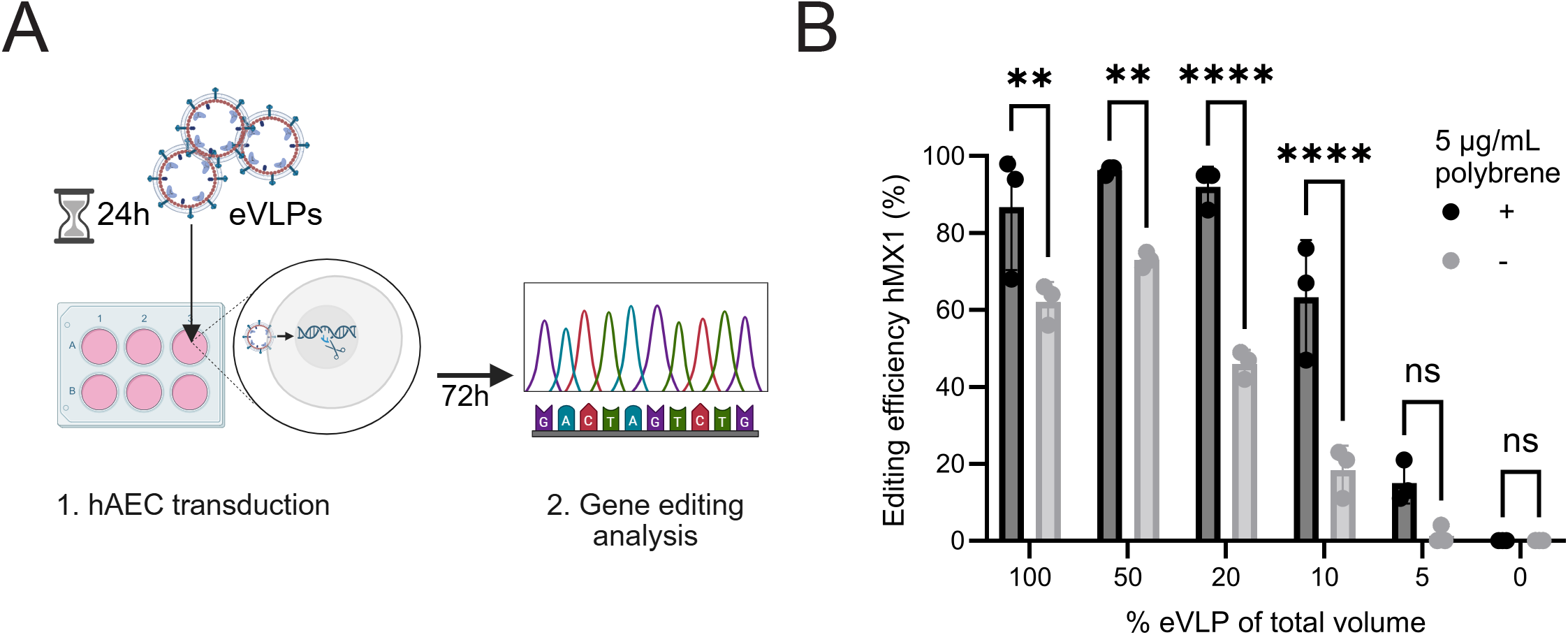
eVLP-mediated genome editing in primary human airway epithelial cells. **(A**) Schematic representation of initial transduction experiment of primary hAECs with eVLPs. Primary hAECs were seeded in a cell culture plate and transduced with supernatant from eVLP producer cells for 24h. At 72h post-transduction, editing efficiency was evaluated by sanger sequencing. Created in BioRender. Stevers, K. M. (2026) https://BioRender.com/2rrw1c0 **(B)** Genomic editing efficiency of hAECs transduced with different volumes of unconcentrated eVLPs targeting *MX1*, as determined by ICE analysis, in the presence (+) or absence (−) of 5 μg/mL polybrene. N=3 biological replicates, bars represent mean with standard deviation. Ns p>0.05, **p ≤ 0.01, **** p ≤ 0.0001 by two-way Anova with Šidák correction for multiple comparisons.

Having demonstrated that eVLPs mediated editing efficiencies approaching 100%, we next set out to develop a workflow that maximized gene-editing efficiency while preserving cell viability and plasticity (Figure 2A). In this workflow, low-passage (<2) primary human airway epithelial cells (hAECs), isolated from healthy bronchial tissue and cryopreserved, are thawed in the presence of the CEPT cocktail, a small-molecule mix reported to reduce stress and improve viability in primary cells (50). Supplementation of CEPT enables transduction of undifferentiated hAECs 24h post-thawing without an initial expansion step, thereby reducing the number of passages required for genetic modification. To minimize potential adverse effects of transduction, hAECs are exposed to eVLPs for only 6h, and, following a medium exchange, editing efficiency is assessed 72h post-transduction.

**Figure 2:**
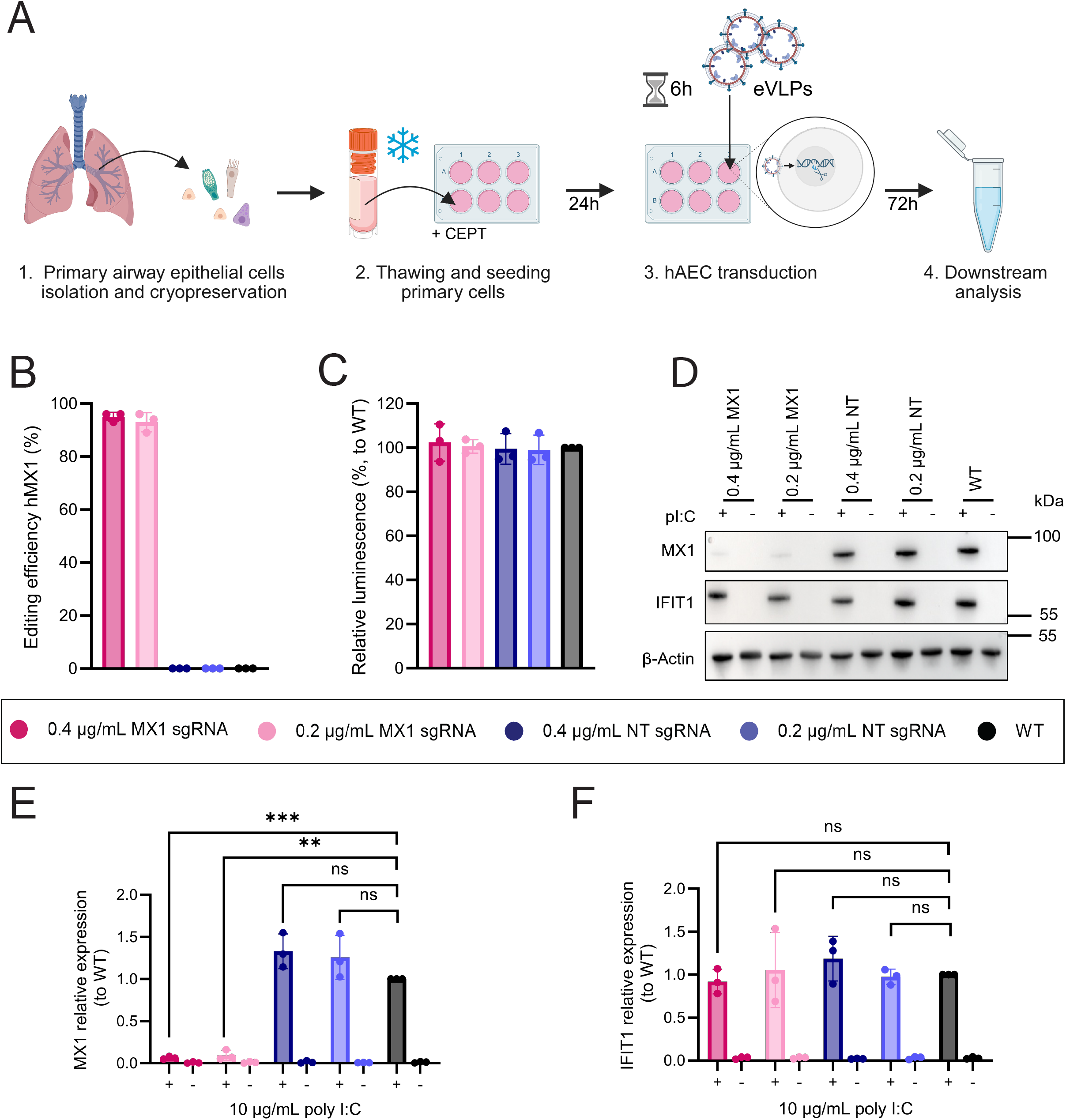
Efficient knockout of MX1 in primary hAEC using eVLPs. **(A)** Schematic representation of the workflow for eVLP-mediated editing of human primary airway epithelial cells (hAECs). hAECs are isolated from tracheobronchial tissue resections and cryopreserved. Cryopreserved cells are thawed in the presence of CEPT (Chroman 1, Emricasan, Polyamine, and trans-ISRIB) small-molecule cocktail to promote post-thawing viability and directly seeded in a cell culture plate. 24h post-seeding, the cells are transduced with eVLPs for 6h. Downstream analyses are performed at least 72h post-transduction. Created in BioRender. Stevers, K. M. (2026) https://BioRender.com/1i576pl **(B)** Editing efficiency of hAECs transduced with 0.4 or 0.2 μg/mL of concentrated eVLPs targeting *MX1*, or non-targeting (NT) eVLPs, compared to untransduced, wildtype (WT) hAECs as determined by ICE analysis. N=3 independent biological replicates, bars represent mean with standard deviation. **(C)** Relative cell viability of hAECs 72 h post-eVLP transduction, as determined by a luminescence-based cell viability assay. N=3 biological replicates, each of which is the average of 4 technical replicates. Bars represent mean with standard deviation. **(D)** Western blot analysis of hAECs transduced with 0.4 or 0.2 μg/mL eVLPs targeting *MX1*, or non-targeting (NT) eVLPs, compared to untransduced, wildtype (WT) hAECs, showing MX1, IFIT1 and β-Actin expression with (+) and without (−) 10 μg/mL poly I:C stimulation (pI:C). **(E-F)** Quantification of western blot protein expression of MX1 **(E)** and IFIT1 **(F).** N=3 independent biological replicates, bars represent mean with standard deviation. Ns p>0.05, **p ≤ 0.01, *** p ≤ 0.001 by Welch t-test with Bonferroni-Dunn correction for multiple comparisons.

To further optimize the system, we sought to reduce the exposure of primary cells to residual contaminants originating from the eVLP-producing cells. To achieve this, eVLP-containing supernatants were concentrated 25-50× by ultrafiltration using a 100 kDa cutoff membrane. Following, eVLP stocks were quantified using a Cas9 ELISA, enabling standardized dosing and reducing the impact of batch-to-batch variation (Figure S1). Using this optimized eVLP production and transduction workflow, we evaluated the editing efficiency and toxicity of concentrated eVLPs at two concentrations (0.4 and 0.2 μg/mL), corresponding to approximately 4× and 2× the 100% volume dose used in Figure 1B, respectively. Both concentrations resulted in >95% editing efficiency at the *MX1* locus (Figure 2B). To assess whether eVLP transduction affected cell viability, we compared hAECs treated with *MX1*-targeting or non-targeting (NT) eVLPs to wild-type, untransduced controls. No measurable differences in viability were observed between conditions (Figure 2C), indicating that eVLP treatment does not adversely affect hAEC viability. To determine editing efficiency at the protein level, interferon-stimulated gene (ISG) expression was induced with the toll-like receptor 3 agonist poly I:C. Protein levels of MX1 and IFIT1, the latter used here as a proxy for general ISG upregulation, were analyzed 24h post-stimulation. Western blot quantification showed that transduction with 0.4 μg/mL eVLPs reduced MX1 protein levels by more than 90%, whereas 0.2 μg/mL eVLPs resulted in an approximately 85% reduction, consistent with the editing efficiencies observed at the genomic level (Figure 2D, E). In contrast, IFIT1 expression remained comparable across all conditions (Figure 2D, F), suggesting that eVLP transduction does not impair ISG induction following poly I:C stimulation. Together, these results demonstrate that this optimized genetic modification workflow enables efficient MX1 knockout (KO) while being well tolerated in primary hAECs.

### eVLP-mediated editing of airway progenitor cells preserves epithelial barrier function and differentiation at the air-liquid interface

Having determined that eVLPs mediated a near-100% MX1 knockout in hAECs, we next evaluated whether the edited cells could be expanded, form a functional epithelial barrier, and differentiate into a multicellular epithelium. Undifferentiated primary airway epithelial cells cultured in 2D conditions, such as during our transduction workflow, have limited regenerative potential and are therefore unsuitable for long-term expansion and cryopreservation of genetically modified hAECs produced on a small scale (33). Alternatively, culturing hAECs in an extracellular matrix within self-organizing 3D airway organoids overcomes these limitations for expansion, but the airway epithelium is oriented inwards, complicating viral inoculation at the apical surface (26,27,51). Therefore, genetically modified hAECs were first expanded as airway organoids, then dissociated into single cells, seeded on semi-permeable inserts, and differentiated at the air-liquid interface for 4-5 weeks, thereby allowing easy access to the apical surface (Figure 3A). We chose to expand and differentiate MX1 KO cells from 3 independent donors, generated with 0.4 μg/mL MX1-targeting eVLPs (hereafter referred to as MX1 KO), together with corresponding NT controls (NT) and naive WT cells (WT).

**Figure 3:**
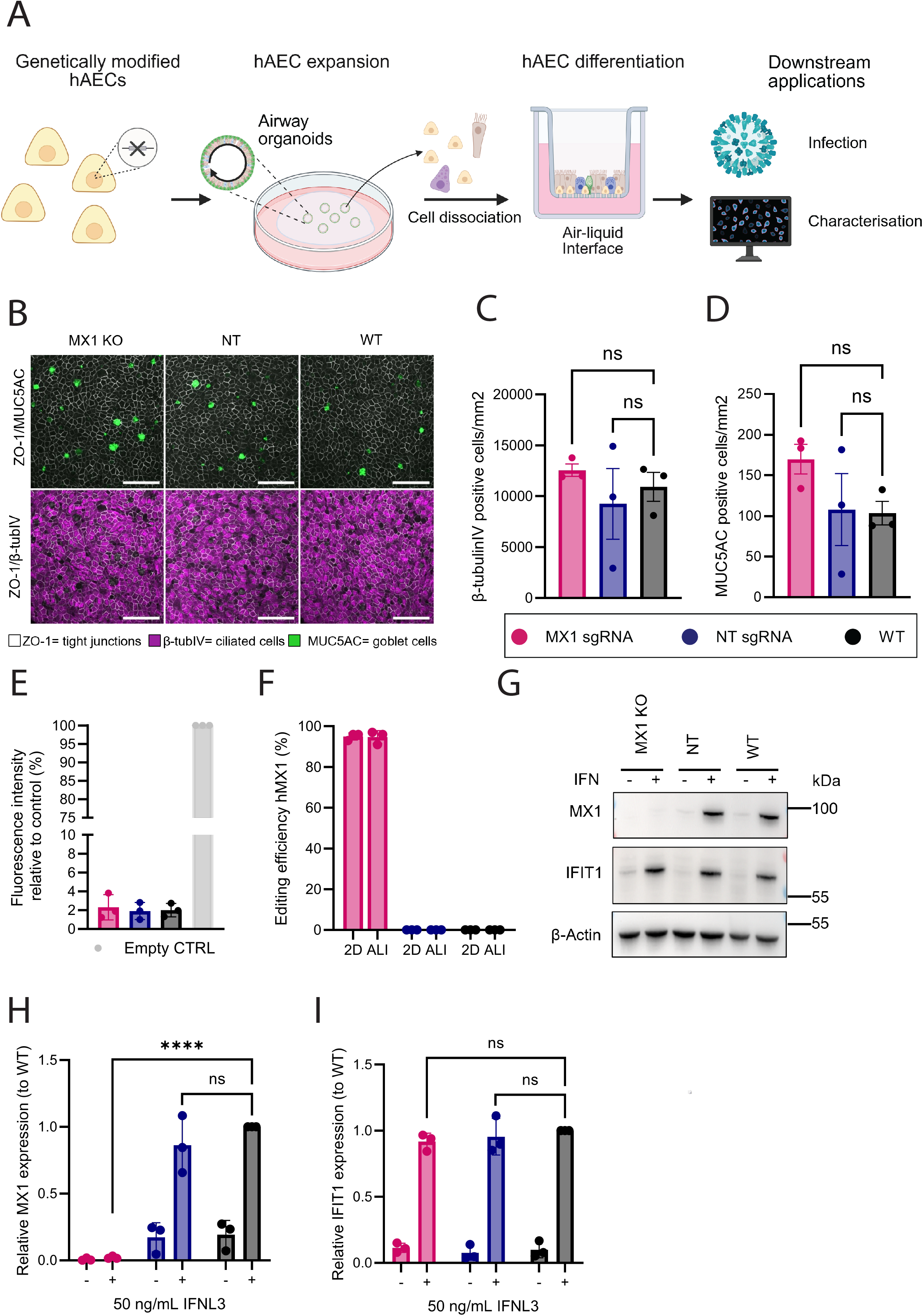
eVLP-mediated editing of airway progenitor cells preserves epithelial barrier function and differentiation at the air-liquid interface. **(A)** Schematic representation of the workflow for expansion and differentiation of genetically modified hAECs. Post transduction, edited cells are expanded as 3D airway organoids (AOs) embedded in a Matrigel extracellular matrix. Following expansion, AOs are dissociated to single cells and seeded on transwell inserts. After establishing a confluent epithelium, cells are exposed to air, driving their differentiation at the air-liquid interface. The resulting multicellular, pseudostratified epithelium resembles the respiratory epithelium *in vivo*, and can be used for downstream applications such as characterization or infections. Created in BioRender. Stevers, K. M. (2026) https://BioRender.com/dna0isr **(B)** Z-projections of well-differentiated MX1 KO, NT or WT ALI cultures. Inserts were fixed and stained with antibodies against ZO-1 (white), MUC5AC (green) or β-tubulin (magenta). Scale bar = 50 μm. Images are representative of N=3. **(C-D)** Quantification of the absolute number of ciliated cells **(C)** and goblet cells **(D)** in 4-week differentiated MX1 KO, NT or WT ALI cultures. Data represents mean with standard error from 8 images each of 3 independent donors per condition. Ns p>0.05 by one-way Anova with Šidák correction for multiple comparisons**. (E)** Relative fluorescence intensity in the basolateral chamber of MX1 KO, NT or WT ALI cultures after 3h incubation with FITC-labeled dextran. Values are shown as percentage of fluorescence relative to control empty insert (100%). 5% is used as a conservative cut-off value of adequate epithelial integrity. N=3 independent biological replicates, each of which is the average of 3-4 technical replicates. Bars represent mean with standard deviation**. (F)** Genomic editing efficiency of MX1 KO, NT or WT hAECs before (2D) and after (ALI) differentiation at the *MX1* locus as determined by ICE analysis. N=3 independent biological replicates, bars represent mean with standard deviation**. (G)** Western blot analysis of MX1 KO, NT, or WT hAEC-ALI cultures, showing MX1, IFIT1 and β-Actin expression with (+) and without (−) 50 ng/mL IFNL3 stimulation (IFN). **(H-I)** Quantification of western blot protein expression of MX1 **(H)** and IFIT1 **(I)** relative to expression in WT hAEC-ALI cultures stimulated with IFNL3. N=3 independent biological replicates, bars represent mean with standard deviation. Ns p>0.05, **** p ≤ 0.0001 by Welch t-test with Bonferroni-Dunn correction for multiple comparisons.

The distinctive mucociliary differentiation and strong epithelial barrier function of hAECs cultured at ALI are essential for recapitulating virus-host interactions and must be preserved when evaluating the effects of gene knockouts to avoid experimental biases arising from altered pathogen access and host responses. Thus, to assess mucociliary differentiation, we employed immunofluorescence staining with the cell markers β-tubulin IV to visualize ciliated cells, MUC5AC to visualize goblet cells, and ZO-1 to visualize tight junctions separating cells (Figure 3B). We also applied unbiased, quantitative image analysis to determine the number of ciliated and goblet cells in MX1 KO, NT, or WT cultures (Figure 3C-D). This analysis showed that MX1 KO, NT, or WT cultures did not differ significantly in the number of MUC5AC- and β-tubulin IV-positive cells, indicating that differentiation was comparable across genetically modified and control cultures. To determine whether epithelial barrier function was sustained in genetically modified cultures, we assessed paracellular permeability using a FITC-labeled dextran permeability assay. The MX1 KO, NT, or WT cultures displayed comparable permeability below 5%, indicating strong barrier function independent of treatment (Figure 3E).

We next investigated the stability of the MX1 knockout during expansion and differentiation; to exclude the possibility that the MX1 knockout cells were negatively selected during this phase. On the genomic level, we observed comparably high levels of gene editing (>95%) before and after differentiation (Figure 3F). To assess the reduction in MX1 protein expression in the differentiated epithelial cells, canonical ISG expression was induced by adding 50 ng/mL of IFNL3 to the basolateral cell culture medium and incubated with the cells for 20h. Following stimulation, we assessed MX1 and IFIT1 protein expression in the differentiated airway epithelium by western blot analysis. Consistent with findings in undifferentiated cultures, MX1 protein was effectively depleted in MX1 knockout cultures but not in control conditions, demonstrating that the knockout phenotype was maintained throughout expansion and differentiation (Figure 3G–H). In contrast, IFIT1 expression was comparable across all conditions, indicating that the overall ISG response remained intact (Figure 3G–I). Collectively, these data establish eVLP-mediated editing as a robust approach for generating KO hAECs while fully preserving the epithelial architecture and mucociliary differentiation required for physiologically relevant ALI cultures.

### Interferon-stimulated MX1 controls the replication of animal-derived influenza A viruses in hAEC ALI cultures

With well-differentiated MX1 KO hAEC ALI cultures established, we subsequently evaluated how MX1 expression in the airway epithelium influenced replication kinetics of IAVs from human or animal origin. As a model virus for human-adapted IAVs, we used the recombinant A/Hamburg/4/09 (H1N1pdm09), whose sequence was isolated from a patient in Germany during the 2009 H1N1 pandemic (52). As models for animal-derived IAVs, we used the recombinant A/Anhui/1/2013 (H7N9) and A/bovine/Texas/2024 (H5N1) virus strains. The sequence of H7N9 was isolated from a patient in China in 2013 (53). The sequence of the H5N1 highly pathogenic avian influenza virus (HPAIV) was isolated from a dairy cow in the United States in 2024, and belongs to clade 2.3.4.4b. genotype B3.13, currently circulating in cattle with reported spillover to humans (54,55).

In immortalized cell lines and in a mouse model overexpressing human MX1, H1N1pdm09 displayed limited sensitivity to MX1, while both H7N9 and H5N1 were significantly restricted (16,56,57). Further evidence in cell lines suggests that MX1 restriction *in vitro* requires IFN pre-stimulation (13,16,58). To determine whether endogenous MX1 expression in the airway epithelium can restrict these IAVs to a similar degree, we pre-stimulated our cultures for 20h with 50 ng/mL of IFNL3, which potently upregulated MX1 expression in NT and WT but not in MX1 KO conditions (Figure 3H). Similarly, unstimulated control conditions, but not MX1 KO cultures, exhibited low baseline MX1 expression (Figure 3H). IFN-stimulated or mock-stimulated MX1 KO, NT, or WT cultures of 3 donors were subsequently inoculated with 10,000 TCID_50_ of each respective virus, and replication kinetics were monitored over 72h. We compared virus replication in IFN-treated and mock conditions to determine the relative contribution of MX1 in the IFN-mediated restriction of IAVs. Each virus replicated to high titers (7-8 log10 TCID_50_ units) under non-stimulated conditions, indicating efficient viral replication (Figure 4A-C). H1N1pdm09 replicated to higher viral titers at earlier timepoints than H5N1 and H7N9. Infections peaked at 48h (H1N1pdm09 and H7N9) or 72h post-infection (H5N1). We did not observe a difference in the replication kinetics of non-stimulated MX1 KO, NT or WT cultures inoculated with H1N1pdm09, H7N9, or H5N1 (Figure 4A-C), suggesting that basal expression levels of MX1 and MX1 expression upregulated after infection are not sufficient to significantly inhibit virus replication. In IFN-stimulated cells, H1N1pdm09 replication in the control cells (NT and WT) was reduced by 2 log10 at the peak of infection (48h) compared to non-stimulated conditions. In contrast, virus production in the MX1 KO hAEC ALI cultures was reduced by only 1 log10, (Figure 4A), suggesting a moderate, IFN-dependent sensitivity of H1N1pdm09 to MX1.

**Figure 4:**
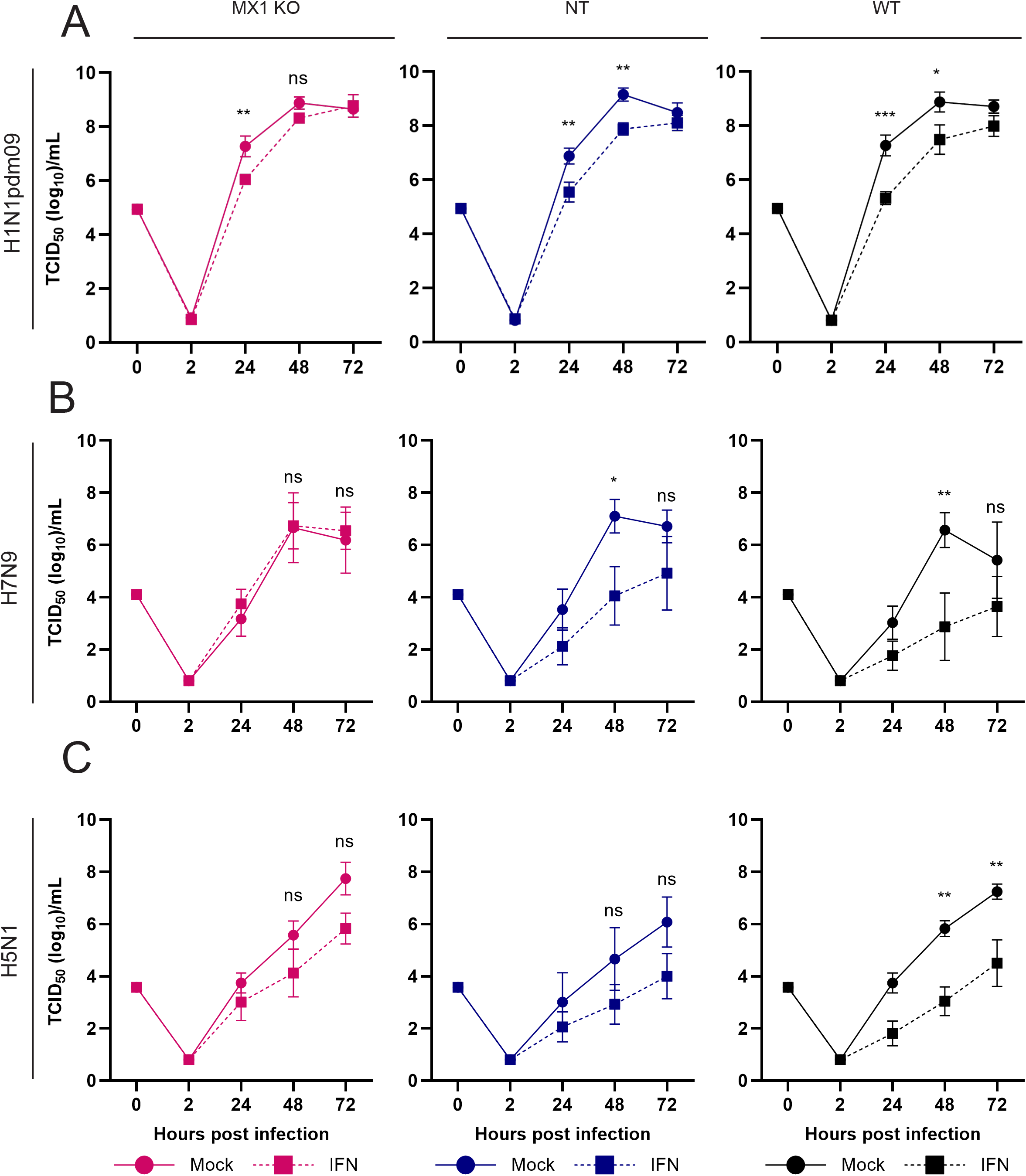
Interferon-stimulated MX1 restricts the replication of influenza A viruses in hAEC ALI cultures. Replication kinetics of H1N1pdm09 **(A)**, H7N9 **(B)** or H5N1 **(C)** IAV viruses as determined by TCID50 on MX1 KO, NT or WT cultures depicted on a log10 scale. Cultures were pretreated for 20h before infection with 0 (mock), 50 (IFN) ng/mL recombinant IFNL3. N=3-6 biological replicates, bars represent mean with standard error. Ns p>0.05, * P ≤ 0.05, **P ≤ 0.01, *** P ≤ 0.001 by two-way Anova with Šidák correction for multiple comparisons.

In contrast, H7N9 replication was reduced by 3 log10 at the peak of viral replication (48h) following IFN pre-stimulation of NT and WT hAEC ALI cultures. However, this reduction in replication was completely reversed in MX1 KO hAEC ALI cultures (Figure 4B). These findings indicate that the IFN system in hAEC ALI cultures is significantly less effective at restricting H7N9 IAV replication in the absence of MX1, suggesting that MX1 is the main interferon-induced factor controlling H7N9 infection in the primary airway epithelium. In contrast, H5N1 replication at the infection peak (72h) was reduced by 3–4 log10 in NT and WT controls, whereas this reduction was only ∼2 log10 in MX1 KO cultures (Figure 4C). This indicates that MX1 contributes to, but is not solely responsible, for IFN-mediated restriction of H5N1 under these conditions.

Because bovine H5N1 was relatively sensitive to interferon λ stimulation in these experiments, consistent with previous reports (59), we reasoned that the strong IFN response induced by 50 ng/mL IFNL3 might saturate the ISG response and potentially mask the effect of the MX1 KO. To test this, MX1 KO, NT, and WT hAEC ALI cultures were pre-stimulated with a lower concentration of IFNL3 (10 ng/mL) prior to H5N1 infection. We first confirmed by Western blot that these less stringent conditions still induced robust MX1 expression in WT cells. Similar to the stimulation with 50 ng/mL IFNL3, treatment with 10 ng/mL IFNL3 for 20h resulted in strong upregulation of MX1 protein (Figure 5A). Upon infection, pre-stimulation with 10 ng/mL IFNL3 significantly reduced H5N1 replication in NT and WT cultures compared with unstimulated controls (Figure 5B). In contrast, viral replication in MX1 KO cultures was restored to levels approaching those observed in unstimulated cells (Figure 5B). This suggests that IFN-induced MX1 can strongly inhibit the replication of bovine H5N1 in hAEC ALI cultures, but loss of MX1 can likely be compensated at higher IFN concentrations with other ISGs.

**Figure 5:**
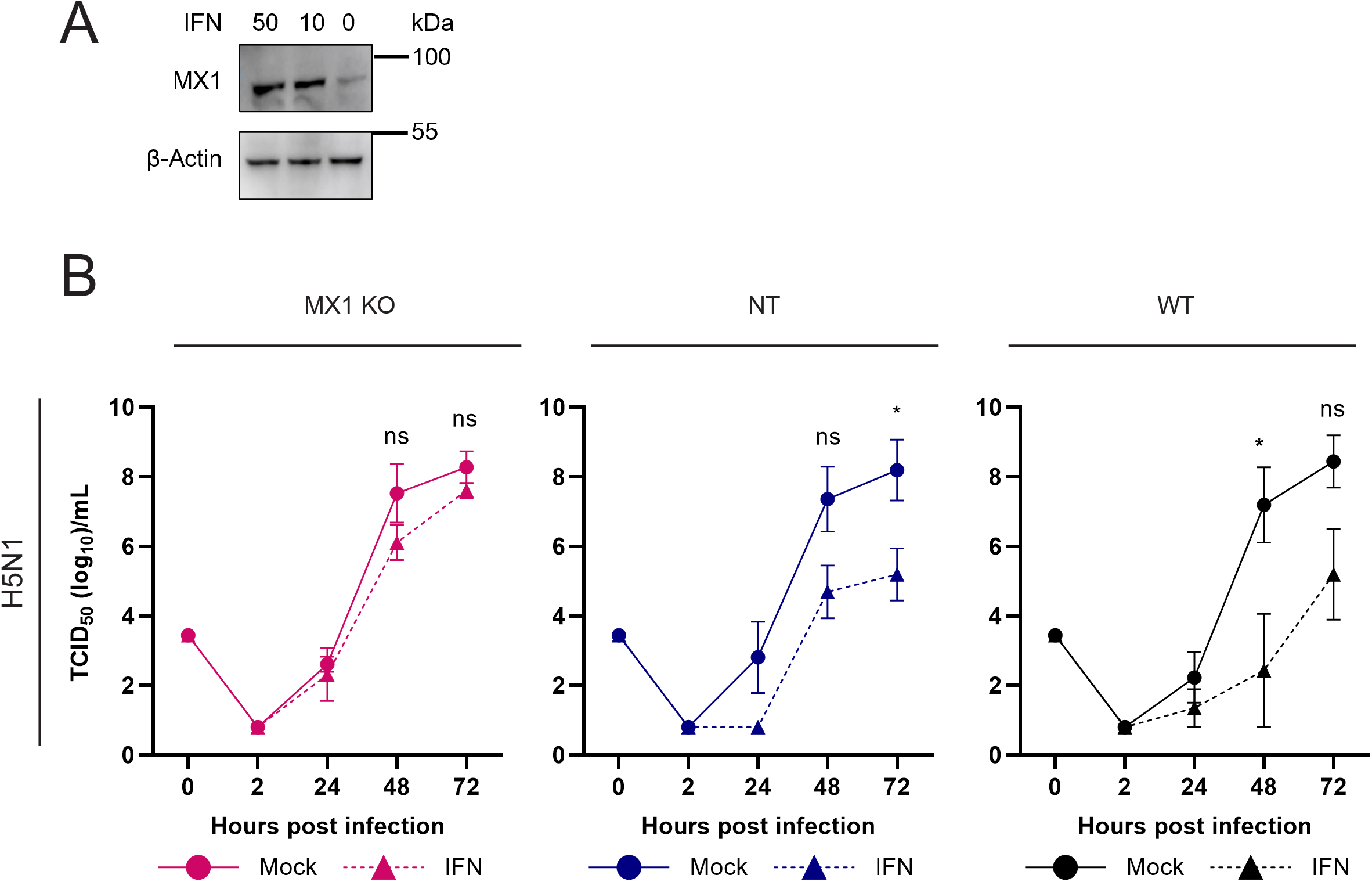
Interferon-stimulated MX1 restricts the replication of bovine H5N1 influenza A virus in hAEC ALI cultures. **(A)** Western blot analysis of WT hAEC-ALI cultures post differentiation at ALI, showing MX1, and β-Actin expression with 0, 10 or 50 ng/mL IFNL3 stimulation (IFN). **(B)** Replication kinetics of H5N1 IAV virus as determined by TCID50 on MX1 KO, NT or WT cultures depicted on a log10 scale. Cultures were pretreated for 20h before infection with 0 (mock), 10 (IFN) ng/mL recombinant IFNL3. N=3 biological replicates, bars represent mean with standard error. Ns p>0.05, *P ≤ 0.05, **P ≤ 0.01 by two-way Anova with Šidák correction for multiple comparisons.

## Discussion

The interferon-stimulated gene MX1 has been widely recognized as a critical ISG and proposed to act as a species barrier for zoonotic IAVs, but this concept has largely been supported by studies using MX1 overexpression systems in immortalized cell lines or transgenic animal models (60). Consequently, the contribution of endogenous MX1 at the primary site of infection in humans, the airway epithelium, has remained incompletely understood. By developing an efficient gene-editing pipeline for primary human airway epithelial cells and generating well-differentiated MX1 knockout cultures, we examined the antiviral defense function of MX1 against IAVs in a physiologically relevant model.

Functional genetic studies in the primary human airway epithelium have historically been constrained by suboptimal gene editing tools and the restricted proliferative capacity of hAECs (33,61). These technical barriers have pushed the field toward alternative strategies for studying host proteins involved in viral infection and replication in airway epithelial cells, such as pharmacological inhibition, antibody-mediated protein blockade or using genetically engineered intestinal organoids as an alternative model (62–64). To address these limitations, we established a novel gene-editing pipeline leveraging the recently developed engineered virus-like particles, which achieved greater than 90% reduction in protein expression in hAECs without the need for selection or clonal expansion. This editing efficiency was achieved while preserving epithelial barrier composition and function after differentiation at the air-liquid interface. Considering eVLPs can be loaded with gRNAs targeting virtually all genomic loci, the complete eVLP-based hAEC editing pipeline provides a versatile platform for generating hAEC ALI cultures perturbing the expression of any host protein of interest. This enables the direct and systematic identification of viral host factor dependencies, as well as the genetic dissection of other human disease mechanisms in a physiologically relevant model system. We have since applied this eVLP-editing workflow to knock out a second innate immune factor with comparable efficiency, underscoring the robustness, reproducibility, and broader applicability of this platform (65).

While multiple studies indicate that electroporation of Cas9-sgRNA RNPs or lentiviral delivery of the Cas9 gene-editing machinery can be as effective as eVLP-based delivery for hAEC gene editing (34,35,66,67), eVLPs offer key advantages. Electroporation relies on membrane permeabilization by high-voltage pulses for delivery, which can induce significant cellular stress (39). In contrast, eVLPs protect cellular integrity by using endogenous endocytosis pathways for entry (42,45), and, as observed in the current study, are well tolerated (Figure 3). Unlike traditional lentiviral vectors, eVLPs lack genetic material that can integrate into the host genome and could cause aberrant gene expression or oncogenesis (41,68). Moreover, by transiently expressing the RNP complex, eVLPs reduce the likelihood of off-target effects associated with prolonged Cas9 expression in lentivirus-transduced cells (40,45). Together, these features make eVLPs an attractive platform for minimally disruptive genome editing of primary hAECs.

By comparing the replication kinetics of IAV in MX1 KO and control hAEC ALI cultures, this study provides, for the first time, direct evidence for the role of endogenous MX1 in restricting zoonotic IAV infection in the human airway epithelium. While loss of MX1 in ALI cultures enhanced replication of all tested IAV strains, the magnitude varied markedly among viral strains of human or animal origin. The MX1 deficiency restored the replication of animal-derived H7N9 and H5N1 viruses in interferon-restricted conditions, albeit for H5N1 only at lower IFN concentrations, while the replication kinetics of human-adapted H1N1pdm09 were affected to a much smaller extent. This suggests that MX1 is a key host factor capable of significantly limiting the replication of animal IAVs in the human respiratory tract. This is in line with, and adds to, core conclusions drawn from landmark studies that positioned MX1 as a major hurdle to cross-species transmission of IAVs to humans (16,18).

The reduced sensitivity of human-adapted IAVs to MX1-restriction compared to those of animal origin is thought to originate from adaptations in the viral nucleoprotein (NP) acquired during sustained transmission between humans or in mammalian intermediate hosts, such as pigs, that express antivirally active forms of MX1 (16,17,20). Site-directed mutagenesis studies have identified several amino acids in the NP of human-adapted H1N1pdm09 virus that contribute to MX1 resistance (17). So far, no amino acids in the H5N1 NP have been demonstrated to confer resistance to human MX1, although bovine MX1 has antiviral properties (56,57). In contrast, H7N9 contains a residue that confers partial MX1 resistance in immortalized cell lines overexpressing MX1 (15,21). This was insufficient to overcome the significant MX1 restriction in the airway epithelium, suggesting that known MX1-resistance-associated residues in NP do not fully predict the phenotype observed in hAECs. Therefore, it remains to be determined whether the MX1-escape mutations identified in cell lines similarly confer MX1 resistance in the human airway epithelium.

The observation that the effect of MX1 KO on IAV replication was evident only with IFN-pretreatment indicated that baseline MX1 expression, as well as MX1 expression induced by infection, is insufficient to significantly restrict viral replication. It is well established that IAVs suppress the host innate immune response through the actions of several viral proteins, most prominently the non-structural protein 1 (NS1) (69). H7N9 and H5N1 are known to elicit a relatively mild innate immune response in human airway epithelial cells, suggesting strong innate immune evasion mechanisms (59,70). This might explain why H5N1 and H7N9, although sensitive to antiviral restriction by MX1, can replicate to high titers in IFN non-stimulated hAEC ALI cultures.

Overcoming MX1-restriction is considered a critical step in the adaptation of emerging IAVs to humans (60). Therefore, understanding which IAVs currently circulating in animal reservoirs have already crossed this boundary is important for biological risk assessment and pandemic preparedness. Established *in vitro* systems based on immortalized cell lines can facilitate rapid assessment of MX1 sensitivity but predominantly rely on non-physiological MX1 overexpression, which artificially enhances its anti-influenza restriction ability, whereas hAEC ALI cultures more accurately reflect physiological MX1 expression and the resulting antiviral restriction. Consistently, the degree of antiviral restriction by MX1 in the hAEC ALI model was smaller for individual strains and between strains compared with these models (15,19,56,57).

Mouse models that express human MX1 are valuable to recapitulate the effect of this host factor on viral virulence and pathophysiology of emerging IAVs *in vivo*, but are limited by inherent species-specific differences between mice and humans that can alter influenza virus infection dynamics, such as the lack of α2,6 sialic acid receptor expression in the airways of mice (71,72), and raise ethical concerns. The MX1 KO hAEC ALI model recapitulates the polarized, pseudostratified epithelium and mucociliary differentiation of the human airway epithelium, the primary site of influenza virus infection in humans. Previous studies have shown primary airway epithelial cell cultures also mimic the transcriptome, sialic acid distribution, influenza virus tropism, and replication kinetics of human airway epithelial tissue (27,30,73). Thus, by comprising key features of IAV infection and endogenous antiviral restriction in the human airway epithelium, the MX1 KO hAEC ALI cultures provide a complementary, physiologically relevant model for evaluating the MX1 sensitivity of emerging IAVs.

In conclusion, eVLP-mediated genome editing provides an efficient and well-tolerated approach for functional genetic studies in differentiated primary human airway epithelium. Using this platform, we demonstrate that IFN-induced MX1 makes a major contribution to the restriction of animal-origin IAVs in the human airway epithelium. These findings suggest that MX1-deficient airway epithelial cultures are a biologically relevant model for directly measuring MX1 sensitivity at the primary site of respiratory infection. These primary cell cultures can provide insights that can be combined with virological, genetic and epidemiological evidence to evaluate the degree to which emerging IAVs have acquired features that increase their zoonotic and pandemic potential.

## Material and methods

### Cell lines

The human embryonic kidney cell line 293LTV (Cellbiolabs; LTV-100) was cultured in 293-LTV medium, the Madin-Darby Canine Kidney II cell line (MDCK-II, ATCC, CRL-2936™) was cultured in MDCK2 medium as described previously (74). Media composition can be found in supplementary table S1. Cells were cultured in a humidified incubator at 37°C with 5% CO_2_ and routinely passaged using phosphate buffered saline (PBS) for washing and TrypLE™ Express Enzyme (Gibco™, Cat-N° 12605036) for dissociation.

### Primary airway epithelial cells

Primary human airway epithelial cells (hAEC/hAECs) were derived from tracheobronchial tissue of patients (>18 years old) undergoing pulmonary resection and were isolated and expanded as described previously (28). Tissue material was obtained through the Tissue Bank Bern, in accordance with ethical approval (KEK-BE 1571/2019).

### Airway Organoid expansion

Organoids were established using a protocol and grown in medium adapted from Sachs et al (26). Briefly, primary hAECs were embedded in ice-cold Cultrex Reduced Growth Factor Basement Membrane Extract, Type 2 (BME, R&D systems, #3533-010-02) with a density of 500-1000 cells/μL BME, depending on the donor. Droplets of 30 µL were seeded onto prewarmed 48-well suspension culture plates (Greiner, Cat-N° 677102) and incubated inverted for 15 minutes at 37°C in a humidified incubator with 5% CO_2_ to solidify. Afterwards, 300 μL fresh airway organoid medium (AOM, supplementary table S1) was added to the domes and cells were incubated at 37°C in a humidified incubator with 5% CO_2_. The medium was refreshed every 3-4 days.

### Organoid passaging

Organoids were passaged every 1-2 weeks. For passaging, AOM on the cells was collected and 300 µL of 1 mg/mL Dispase II solution (Sigma-Aldrich, Cat-N° D4693, diluted in DMEM (Gibco, Cat-N°10566016)) was added to each well to dissolve the BME. Cells were incubated for 2 hours at 37°C. After two hours, the cells were harvested by mechanically disrupting the domes with a pipet tip, transferring the Dispase/cell mixture to a fresh 15 mL tube, and centrifuging at 500 × g for 5 minutes. The supernatant was carefully aspirated, and cells were resuspended in TrypLE (100-150 µL per dome) and incubated for 15 minutes at 37°C, with pipetting up and down every 5 minutes. After dissociation to single cells, TrypLE was inactivated by adding the old medium, collected prior to adding the dispase II solution, and cells were centrifuged at 500 x g for 5 minutes. The supernatant was carefully aspirated, and the pellet was resuspended in 0.5-5 mL of pre-AOM (supplementary table S1) supplemented with 5 µM Y-27632 (Tocris, Cat-N°1254). Cells were counted and the necessary number of cells were transferred to a fresh 15 mL tube. Cells were centrifuged at 500 x g for 5 minutes, the supernatant was carefully pipetted off and the pellet was resuspended in ice-cold BME, with a density between 250-750 cells/µL BME, depending on the donor. Cells in BME were seeded and cultured as described above.

### Organoid freezing and thawing

Organoids were routinely frozen at −150°C post-dissociation in Cryostor CS10 (StemCell, Cat-N° 07930) at a density of 1E6 cells/mL, according to the manufacturer’s instructions. Cells were thawed by warming the cryovial in a 37°C water bath, transferring the contents to a 15 mL tube, and adding 4 mL of prewarmed pre-AOM, supplemented with 1x CEPT cocktail (Tocris, Cat-N° 7991) dropwise to the cells. Afterward, cells were centrifuged at 200 x g for 3 minutes at RT, and the supernatant was carefully pipetted off. The cell pellet was resuspended in 1 mL pre-AOM supplemented with 1x CEPT and centrifuged at 200 x g for 3 minutes at RT. The cell pellet was resuspended in BME at a density of 500-1000 cells/μL BME and plated as described above. In the first 2-3 days post-thawing, organoids were grown in AOM supplemented with 1x CEPT, before switching back to regular AOM.

### Air-liquid interface culture

Air-liquid interface (hAEC ALI) cultures were generated using organoid-derived AECs from passages 3 to 7. Cell culture inserts (6.5 mm diameter, CellQuart, Cat-N° 9320414) were coated with 50 μg/mL collagen type IV solution (Sigma, Cat-N° C7521, dissolved in 0.25% acetic acid). A single cell suspension of AECs was prepared as described in organoid passaging, resuspended in epithelial expansion medium (EpiX, Pneumacult Ex Plus medium Stemcell, Cat-N°5040, supplemented with previously described EpiX formulation (75), supplementary table S1) and seeded on coated inserts with a density of approximately 100,000 cells/insert. Cells were expanded on inserts until confluency in an initial liquid-liquid state (3-6 days) in EpiX. Medium was replaced every 2-3 days. Once the cells reached full confluency, the basolateral medium was replaced for complete ALI medium (Pneumacult-ALI medium, Stemcell, Cat-N°5001, supplementary table S1). The apical side was washed with TEER buffer (0.9% NaCl, 1.25 mM CaCl_2_, 10 mM HEPES) and exposed to air to establish ALI. Cells were cultured under ALI conditions for 3-5 weeks until differentiation, which was visually assessed by the presence of beating cilia and mucus production and monitored by light microscopy. Cells were maintained at 37°C in a humidified incubator with 5% CO_2_ and cultured by refreshing the complete ALI medium every 2-3 days. The apical side of the inserts were washed every 2 weeks with TEER buffer by adding 200 μL buffer, incubating for 10 min at 37°C and pipetting up and down for at least 5 times.

### Viruses

The reverse genetic plasmids for influenza A/Hamburg/4/09 (H1N1pdm09) were kindly provided by Prof. Martin Schwemmle, University of Freiburg, Germany, whereas the plasmids for A/Anhui/1/2013 (H7N9), were kindly provided by David Wentworth, J. Craig Venter Institute, Rockville, Maryland, USA, and described previously (19,76). Recombinant viruses were rescued from cocultures of MDCK-II and 293-LTV cells transfected with 1 μg of each genomic plasmid as described previously (74), using lipofectamine LTX (Invitrogen, Cat-N° 15338100) at a 1:2 DNA:LTX ratio, according to the manufacturer’s instructions. Viral supernatant was harvested when >50% CPE was observed and clarified by centrifugation at 1000 x g for 5 minutes. A working stock of all viruses was generated by infecting MDCK-II cells with 500 µL of viral supernatant from the rescue in infection medium (iMEM, supplementary table S1), supplemented with 1 µg/mL bovine pancreas-isolated acetylated trypsin (Sigma-Aldrich, Cat-N° T6763), for H1N1pdm09, with harvesting after 24-48h. Viral supernatant was clarified by centrifugation at 1000 x g for 5 minutes, and virus was aliquoted and frozen at −80°C until use. The Influenza A/bovine/Texas/24-029328-02/2024 (H5N1) was generated as described previously (59).

### Virus titration

The titers of the viral stocks and apical washes were determined by the tissue culture infectious dose 50% (TCID_50_) using MDCK2 cells. 24h before titration, MDCK2 cells were seeded in a 96-well plate at a density of 30,000 cells/well. The next day, the culture medium was aspirated, cells were washed with PBS, and 50 µL iMEM was added to each well. Each sample was serially diluted 10-fold in iMEM, and 50 µL of each dilution was added to cells in 6 technical replicates. For H1N1pdm09, iMEM was supplemented in each step with 1 µg/mL of trypsin. Cells were incubated with virus for 3 days at 37°C in a humidified incubator with 5% CO_2._ After 3 days of incubation, the supernatant was aspirated, cells were fixed with 4% formalin, and CPE was visualized with 0.5% crystal violet solution. TCID_50_/mL was calculated based on the Spearman-Kärber formula (77).

### eVLP production

The following plasmids were used for eVLP production. SUPERBLADE5 (a gift from Philippe Mangeot & Théophile Ohlmann & Emiliano Ricci, Addgene #134913, (42)), pMD2.G (Addgene #12259, a gift from Didier Trono), pBS-CMV-gagpol (Addgene #35614, a gift from David Liu, (45)) and pCMV-MMLVgag-3xNES-Cas9 (Addgene #181752, a gift from David Liu, (45)).

eVLPs were generated using a protocol adapted from Banskota et al (45). The gRNA targeting MX1 (5’ - 3’: GATATGTGTGATGAGCTCGC) was designed using CHOPCHOP (78), and the non-targeting gRNA sequence (5’ - 3’: AAATGTGAGATCAGAGTAAT) was obtained from Thermo Fisher Scientific (Cat-N° A35526). The gRNAs were cloned into the SUPERBLADE5 vector using the BsmBI restriction site. For eVLP production, 293-LTV cells were seeded on poly-D-lysine-coated plates (Sigma-Aldrich Cat. No. P0899) at a density of 6.6E6 cells/T75 flask. The eVLPs were produced by transfecting 293-LTV cells with the SUPERBLADE5-gRNA, pBS-CMV-gagpol, pCMV-MMLVgag-3xNES-Cas9, and pMD2.G vector using JetOptimus (Polyplus, Cat-N° 101000051), in a 1:1 reagent:DNA ratio according to the manufacturer’s instructions. Plasmid ratios can be found in Supplementary table S2. The day after transfection, the medium was replaced by EpiX medium. 48-72h post-transfection, the eVLPs were harvested and clarified twice by centrifugation at 1000 x g for 5 minutes. Supernatant containing eVLPs was stored at −80°C for unconcentrated use or concentrated by ultrafiltration.

eVLPs were concentrated using Amicon® Ultra-15 100 kDa MWCO Centrifugal Filter Units (Merck Millipore, Cat-N° UFC910096) at 1500x g for 45 minutes at 4°C. eVLPs were harvested by collecting the concentrate and washing the filter membrane with 200-400 uL EpiX medium. Concentrate and wash were combined, and 0.01% of Pluronic F-68 (Gibco, Cat-N°24040-032) was added to the concentrated eVLPs to reduce adhesion to plastic. eVLPs were aliquoted and stored at −80°C. Generally, eVLPs were used within 3 months of production.

### eVLPs quantification

eVLPs were quantified using the Cas9 ELISA Kit (Cell Biolabs, Cat-N° PRB-5079), according to the manufacturers protocol. Prior to the protocol, concentrated eVLPs were diluted to between 1:100 and 1:1000 in assay diluent (Cell Biolabs Cas9 ELISA kit) and inactivated with 0.5% Triton X-100 at 37°C for 30 minutes. The Cas9 standard protein (Cell Biolabs Cas9 ELISA kit) was serially diluted according to the manufacturer’s protocol and equally treated with 0.5% triton X-100 at 37°C for 30 minutes. Absorbance was measured at 450 nm using the Biotek Cytation 5 Cell Imaging Multimode Reader (Agilent).

### Determination of eVLP dose

To determine the optimal dose for eVLP transduction and evaluate the effect of polybrene addition, primary hAECs were seeded with a density of 30,000 cells/cm2 in collagen I-coated 24-well plates (40 ug/mL, Sigma-Aldrich, Cat-N°C2249). The next day, unconcentrated eVLPs were thawed on ice and diluted in KSFM-EpiX medium (supplementary table S1) to 5, 10, 20, 50 and 100% volume of 500 uL. Medium on cells was aspirated and eVLPs were added to the cells, supplemented with 0 or 5 μg/mL polybrene (Sigma, Cat-N° TR-1003-G). 24h post transduction, inoculum was aspirated, cells were washed once with Hank’s balanced Salt Solution (HBSS, Gibco, Cat-N° 14175095) and fresh KSFM-EpiX medium was added to each well. eVLP-mediated gene editing efficiency was determined 72h post transduction.

### eVLP transduction of primary airway epithelial cells

For all subsequent experiments, Low passage (<2) primary human AECs, were thawed in a 37°C waterbath and transferred to a 15ml falcon tube. 5 mL thawing medium (see supplementary table S1) consisting of EpiX medium supplemented with 1x CEPT cocktail (Tocris, Cat-N° 7991), was added dropwise to the cells. Cells were centrifuged at 200 x g for 3 minutes at RT, and the supernatant was carefully pipetted off. The cell pellet was resuspended in 1 mL thawing medium, counted using a Countess II Automated cell counter (Invitrogen), and seeded with a density of 30,000 cells/cm2 in collagen I-coated 24-well plates. The next day, eVLPs were thawed on ice and diluted to 0.2 and 0.4 μg/mL Cas9 in EpiX medium to a total volume of 500 μL per 24-well. Medium on the cells was aspirated and diluted eVLPs were added to the cells in the presence of 5 μg/mL polybrene. 6h post transduction, the inoculum was aspirated, the cells were washed once with HBSS and fresh EpiX medium was added to each well and incubated for 72h.

At 72h post-transduction, the supernatant was aspirated, and the cell monolayer was washed with HBSS. Cells were dissociated with TrypLE, harvested, and washed with pre-AOM. The cells were centrifuged at 500 x g for 5 minutes, and the supernatant was carefully pipetted off. Transduced cells were harvested to determine gene-editing efficiency, reseeded in EpiX medium for protein expression analysis, or embedded in BME and expanded as organoids as described in “organoid expansion”.

### Determination of eVLP-mediated gene-editing efficiency

The eVLP editing efficiency was determined 72h post transduction and 4 weeks post differentiation at ALI by lysing the cells using the DirectPCR Lysis Reagent (Cell) (Viagen, Cat-N°301-C) supplemented with 200 µg/mL Proteinase K (Promega, Cat-N° MC5005), according to the manufacturer’s protocol. A ∼700bp fragment surrounding the target site was specifically amplified with GoTaq G2 Hotstart Green Master Mix (Promega, Cat-N°M7423) using the lysed cells as template. Primer sequences are forward 5’-3’: GACCAAGTCTATAGTCAGACAC and reverse 5’-3’: CCACAGAGAGCTTTTACAAAG. Correct PCR fragment size was verified using a 1.5% agarose gel, and products were purified using the E.Z.N.A. cycle pure kit (Omega Bio-tek, Cat-N° D6492-02), according to the manufacturer’s protocol. Purified amplicons were subjected to Sanger sequencing (Microsynth, Balgach, Switzerland). Sequencing results were analyzed for indel frequency using the Inference of CRISPR Edits webtool (ICE CRISPR Analysis. 2025. v3.0. EditCo Bio, (79)), which was utilized as a proxy for editing efficiency in this study.

### Western blot

Protein expression levels were determined by western blot. To induce ISG expression before differentiation, transduced cells were stimulated with 10 μg/mL poly I:C (HMW, Invivogen, Cat-N° rl-pic) diluted in EpiX medium 6 days post transduction. After 24h stimulation, cells were harvested using M-PER Mammalian Protein Extraction Reagent (Thermo Fisher Scientific, Cat-N° 78501), supplied with 1x Halt™ Protease Inhibitor Cocktail (Thermo Fisher Scientific, Cat-N° 78429). Post-differentiation at ALI, cells were basolaterally stimulated with 50 ng/mL recombinant interferon lambda 3 (IFNL3, (80)) diluted in complete ALI medium for 20h, followed by membrane excision and lysis with M-PER supplemented with 1x Halt according to the manufacturer’s protocol. The lysates were quantified using the Qubit Protein BR Assay kit (Invitrogen, Cat-N° A50668) according to the manufacturer’s protocol. Equal amounts of protein were mixed with a final concentration of 1× Lämmli-buffer (5 mM Tris pH 6.8, 0.8% glycerol, 0.25% SDS, 0.5% β-mercaptoethanol, and 0.005% bromophenol blue), boiled for 5 minutes at 95 °C, and cooled on ice. Samples were loaded on a a 4–12% gradient SurePAGE, Bis-Tris precast gel (Genscript, Cat-N° M00654) and run at 180 V for 45 minutes. Gels were transferred to a nitrocellulose membrane using an eBlot L2 Fast Protein Transfer Device (GenScript). Subsequently, the membrane was blocked using 5% BSA (IgG-free, Protease-free, Jackson ImmunoResearch, Cat-N°001-000-162) in TBST (20 mM Tris, 500 mM NaCl, pH7.4, 0.05% Tween-20), followed by overnight incubation with the primary antibody diluted in TBST with 0.5% BSA at 4°C. For detection, the membrane was incubated for 1h with a secondary HRP-conjugated antibody for 1h at room temperature. Next, the membranes were stripped twice for 10 minutes using stripping solution (0.2 M Glycine, 0.1% SDS, 1% Tween20, pH 2.2), followed by reblocking and incubation with a loading control antibody (anti-β-actin HRP-conjugated). Bands were visualized using the WesternBright Quantum detection kit (Advansta, Cat-N°K-12042-D10) and imaged using the Fusion FX (Vilber). All antibodies and dilutions can be found in supplementary table S3. Western blots were quantified using the EvolutionCapt Edge software and normalized to β-Actin.

### Infection of ALI cultures

Well-differentiated hAEC ALI cultures were infected at 3-5 weeks post ALI. 3-4 days before infection, the basal cell culture medium was replaced with complete ALI medium without heparin and hydrocortisone (ALI infection medium, supplementary table S1). 20h before infection, cultures were basolaterally pre-stimulated in certain conditions with 50 or 10 ng/mL IFNL3 diluted in ALI infection medium, as indicated in the figure legend. Directly before infection, the basolateral medium containing IFNL3 was removed and replaced with ALI infection medium without IFNL3, and cultures were washed twice apically with TEER buffer. hAEC ALI cultures were apically infected with 10,000 TCID50 of H1N1pdm09, H5N1, or H7N9, diluted in HBSS buffer, for 2h at 37°C. Afterward, viral inoculum was removed, and the apical side was washed three times with TEER buffer. The last wash was collected as the 2h time point. The cultures were incubated at 37°C in a humidified incubator with 5% CO_2_ until the indicated time points. At each time point, progeny virus was collected by applying TEER buffer to the apical side and incubating for 10 minutes prior to collection. Collected virus was diluted 1:1 with viral transport medium (VTM, supplementary table S1) and stored at −80°C until further use.

### Cell viability assay

Cell viability of undifferentiated hAECs was determined at 72h post transduction using the CellTiterglo 2.0 viability assay (Promega, Cat-N° G9241), according to the manufacturer’s protocol. Briefly, primary hAECs were seeded at a density of 10,000 cells/well in black clear bottom 96-well plates (Corning, Cat-N° 3603/10530753). The next day, the cells were transduced with eVLPs (as described under eVLP transduction) for 6h. After 72h, cells were lysed with 200 μL CellTiter-Glo reagent 2x diluted with HBSS. Luminescence was measured using the Biotek Cytation 5 Cell Imaging Multimode Reader (Agilent). For each donor and treatment condition, 4 technical replicates were used.

### FITC Dextran permeability assay

The epithelial integrity of the hAEC ALI cultures was determined after 3-4 weeks of differentiation with a FITC Dextran paracellular permeability assay, as described previously (81). Briefly, the apical side of the inserts was washed with 200 μL TEER buffer to remove mucus and the basolateral medium replaced by fresh complete ALI medium. Fluorescein isothiocyanate–dextran (Sigma Aldrich, Cat-N° FD4-100MG) was diluted to 2 mg/mL in HBSS, applied apically to the inserts (150 μL per insert) and incubated at 37°C in a humidified incubator with 5% CO_2_ for three hours in the dark. After incubation, the basolateral medium was transferred to black clear bottom 96-well plates in triplicates, and absorbance was measured at 485 nm using the Cytation 5 Multimode reader. Relative permeability was measured by normalizing the absorbance of each culture to the fluorescence in the basolateral chamber of an empty insert.

### Immunofluorescence staining

hAEC ALI cultures were fixed with 4% buffered formalin for 20 minutes and processed as described previously (28). Briefly, cells were permeabilized and blocked in confocal buffer (CB, PBS supplemented with 50 mM NH4Cl, 0.1% (w/v) Saponin, and 2% (w/v) bovine serum albumin (Jackson Immunoresearch, Cat-N° 001-000-162)). Cell-type specific markers were detected with antibodies against MUC5AC (Novus Biological, Cat-N°NBP2-15196) for goblet cells, Alexa Fluor® 647 conjugated Anti-beta IV Tubulin antibody [EPR16775] (Abcam, Cat-N° ab204034) for ciliated cells, and Alexa Fluor 594 conjugated ZO-1 Monoclonal Antibody (ZO1-1A12) (Thermofischer Scientific, Cat-N° 339194) for tight junctions. Alexa Fluor 488-labeled donkey anti-rabbit (Jackson ImmunoResearch, Cat-N° 711-545-152) was used as secondary antibody.

All antibodies were diluted in CB and antibody dilutions used in this study can be found in Supplementary table S3. To visualize the nuclei, all samples were counterstained using 4′,6-diamidino-2-phenylindole (DAPI, Thermo Fisher Scientific). The immunostained inserts were placed on slides with DakoFluorescence Mounting Medium (Agilent), covered in ProLong™ Diamond Antifade Mountant (Thermo Fisher Scientific, Cat-N° P36965), overlaid with coverslips and allowed to cure overnight.

Images were acquired on a Nikon Ti2 Cicero spinning disk confocal microscope using a 20× air objective NA=0.8). Nine regions of interest (ROI) were systematically selected across the insert using a cross shaped acquisition pattern, covering both central and peripheral zones. Relative to the ZO-1 as middle plane, a Z-stack with 2 µm increments over 48 µm was acquired.

### Image quantification

Fluorescence microscopy images were acquired as multi-channel z-stacks in nd2-file format and analysed using a Python-based image analysis pipeline developed independently by collaborators at the Institute of Virology and Immunology and the Data Science Lab, University of Bern (https://github.com/quasar1357/ivi-imaging commit: 4b55093). Image analysis was performed in Python 3.11.0 using CellPose (v3, cyto3 model, (82)), scikit-image, NumPy, pandas, and aicsimageio. Raw z-stacks were loaded using AICSImage, and maximum-intensity projections were generated independently for each fluorescence channel along the z-axis.

Cell segmentation was performed on the projected images using the ZO-1 tight-junction signal and the CellPose denoising model (CellposeDenoiseModel; model type: cyto3; restore type: denoise_cyto3) with a cell diameter of 40 pixels. Per-cell fluorescence intensities were subsequently quantified for MUC5AC as a goblet-cell marker (50th percentile intensity within the cell mask to minimize signal crosstalk) and β-tubulin IV as a ciliated-cell marker (mean intensity within the cell mask). Fluorescence intensities were log10-transformed before classification. For each marker, a single threshold was automatically determined using Otsu’s method and applied across all images to classify cells as marker-positive or -negative (83). Cells positive for both markers were excluded. The absolute numbers of MUC5AC-positive goblet cells and β-tubulin IV-positive ciliated cells per mm² were quantified and used for subsequent statistical analyses.

### IAV whole genome sequencing

To determine the whole genome sequence of influenza A viruses used in this study, viral RNA was extracted from samples using the KingFisher Apex automated platform (ThermoFisher Scientific) together with the NucleoMag® VET kit (Macherey–Nagel, Cat-N°744200.4) following the manufacturer’s instructions, or extracted manually using the Quick-RNA Viral Kit (Zymo Research, Cat-N°R1034) according to the manufacturer’s protocol.

Whole genome amplification, Nanopore sequencing (Oxford Nanopore Technology), sequence analysis, were performed as described previously (84,85). Basecalling was performed using dorado (version 0.7.11.2, (86)) with the high-accuracy (HAC) model (version 5.2.0). Reads were filtered using Chopper (version 0.9.2) (87). Genomes were assembled using Iterative Refinement Meta-Assembler (version 1.3.1) (88), sequences were aligned using MAFFT (version 7.2) (89), and alignments were visualized with Aliview (version 1.3.0) (90).

### Statistics

Statistical analyses were performed using GraphPad Prism (version 10.2.1; GraphPad Software) on data from three or more independent biological replicates, as indicated in the figure legends. Statistical tests are specified in the corresponding figure legends. Differences were considered statistically significant at *P* < 0.05.

## Acknowledgements

This research was supported by the Swiss National Science Foundation (Grant No. 310030_207675). We would like to thank Martin Schwemmle and David Wentworth for providing the H1N1pdm09 and H7N9 reverse genetics plasmids, respectively. We also thank Vera Flück and Jenna Kelly from the Institute of Virology and Immunology, Bern, Switzerland, and Roman Schwob from the Data Science Lab, University of Bern, Switzerland, for their contributions to the development, implementation, and application of the image analysis pipeline used in this study. We also extend our gratitude to Laura Graf and Neda Hosseini from the University of Freiburg, Germany, for their assistance with the eVLP system. We are grateful for the advice and support of the biosafety team at the Institute for Infectious Diseases, University of Bern, Switzerland. Furthermore, we are grateful to Sabina Berezowska and Irene Ramos-Centeno, Institute of Pathology, University of Bern, for providing the respiratory tract tissues via the Tissue Bank Bern. Microscopic imaging was performed on equipment supported by the Microscopy Imaging Center (MIC), University of Bern, Switzerland

## Conflicts of Interest

The authors declare no conflict of interest.

---

**Figure S1:**
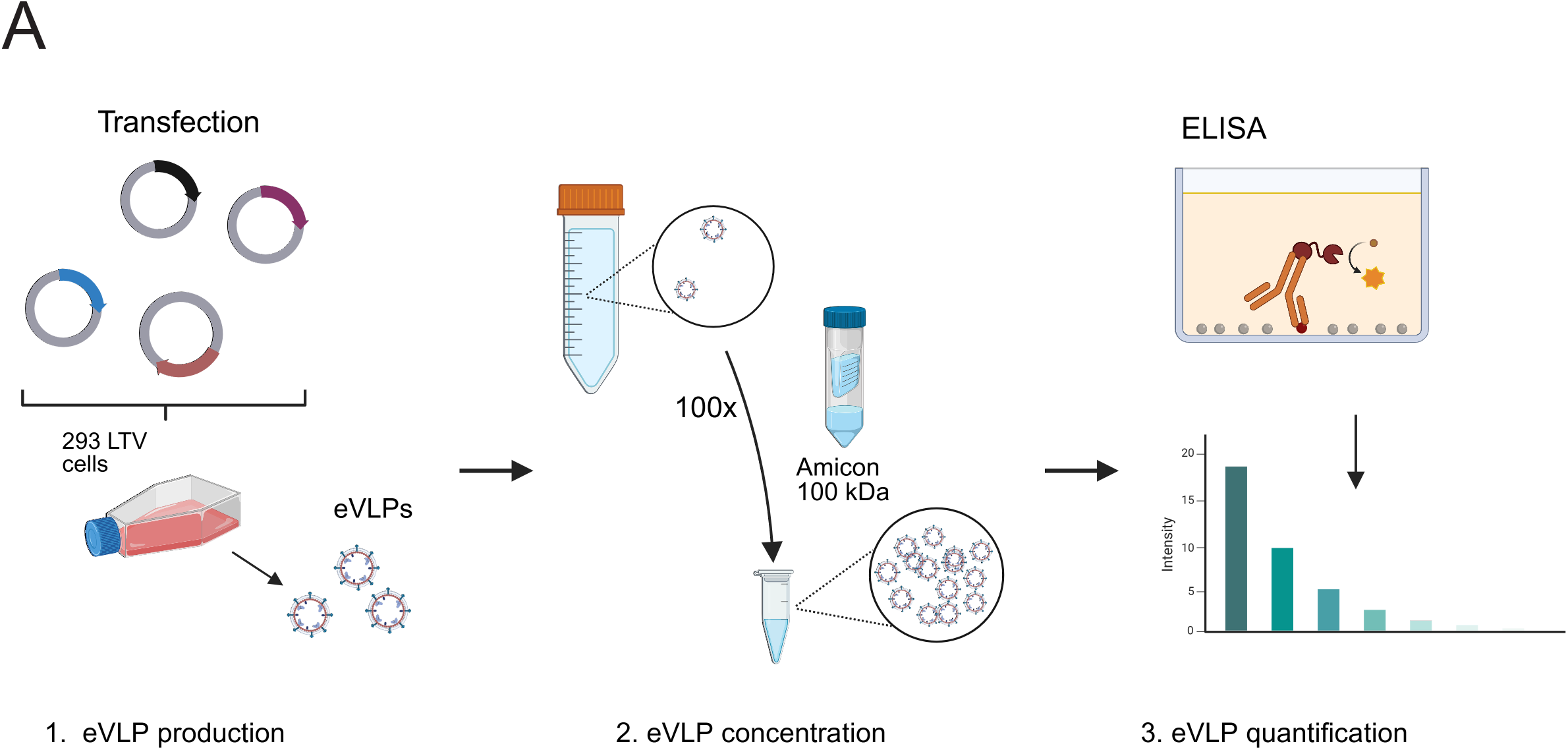
optimized eVLP production, concentration, and quantification workflow. **(A)** Schematic representation of the workflow for production, concentration, and quantification of eVLPs. eVLPs are produced by transfection 293-LTV cells by plasmids encoding the VSV-G glycoprotein, the gRNA, the MMLV structural proteins gag and pol and a plasmid encoding a fusion product of cas9 with the MMLV gag protein. The resulting supernatant containing eVLPs is concentrated using ultracentrifugation with a 100 kDa cut-off filter. These concentrated eVLPs are then quantified using a Cas9 ELISA. Created in BioRender. Stevers, K. M. (2026) https://BioRender.com/cbmcpm3

## Notes

### Competing Interest Statement

The authors have declared no competing interest.

